# Multimodal Analysis Reveals Differential Immuno-Metabolic Features in Lung Squamous Cell Carcinoma and Adenocarcinoma

**DOI:** 10.1101/2021.05.09.443240

**Authors:** Brooks P. Leitner, Shyryn Ospanova, Aray Beisenbayeva, Kevin B. Givechian, Katerina Politi, Rachel J. Perry

## Abstract

**Background:** The relationship between systemic metabolism, immune function, and lung cancer is complex and remains poorly defined. Seemingly paradoxically, overweight and obesity confer an improved response to immune checkpoint inhibition in non-small cell lung cancer (NSCLC); however, it is not known whether excess body weight or adiposity impacts the immunometabolic tumor microenvironment.

**Methods:** Utilizing three complementary National Cancer Institute-funded open-source databases containing ^18^F-fluorodeoxyglucose positron-emission tomography/computed tomography (PET-CT) images for tumor and tissue glucose uptake, adipose tissue and skeletal muscle mass, histology annotated with tumor infiltrating leukocytes, and tumor RNA sequencing, we performed a retrospective cross-sectional analysis to examine phenotypic, metabolic, and genomic intersections of adiposity and tumor immune-metabolism in patients with lung adenocarcinoma (LUAD) versus squamous cell carcinoma (LUSC).

**Results:** Our data reveal distinct immunometabolomic features of LUSC as compared to LUAD: visceral fat content was negatively correlated with both tumor glucose uptake and leukocyte infiltration. Subcutaneous and visceral adiposity conferred different effects on the tumor genetic landscapes in both tumor types. LUSC tumors showed greater gene expression pathways related to pyruvate, glucose, amino acid, and lipid metabolism, in addition to significantly greater ^18^F-FDG uptake compared with LUAD, suggesting deeper metabolic regulation within the LUSC tumor microenvironment.

**Conclusions:** Several immunometabolomic characteristics of LUSC and LUAD differ, including tumor glucose uptake and the associated metabolic pathways in the tumor, as well as the impact of visceral adiposity on tumor metabolism. These data may highlight opportunities to advance mechanistically targeted precision medicine approaches by better understanding the interplay between metabolic, immunologic, and genomic factors in lung cancer treatment.

## Introduction

Lung cancer is one of the few tumor types that is not traditionally associated with metabolic dysregulation. Clinically, it has not been positively associated with excess body weight and obesity by the U.S. Centers for Disease Control^1^, and, interestingly, several reports have suggested that excess body weight may actually reduce the risk and slow the progression of lung cancer^2–6^, particularly in those treated with immune checkpoint inhibitors^7,8^. However, studies of how body composition and metabolism affect lung cancer outcomes have rarely differentiated between the subtypes of lung cancer, and even more rarely have differentiated between the subtypes of NSCLC. This knowledge gap limits the possibilities of developing metabolic strategies to combat lung cancer using a precision medicine approach.

Tumor glucose uptake, most commonly measured by PET-CT with [^18^F]-fluorodeoxyglucose (^18^F-FDG) in humans, has long been utilized as a marker of metabolic activity. Glucose taken up by tumors primarily fuels glycolytic metabolism, which produces glucose- and fructose-6-phosphate and, in turn, generates the cellular building blocks (nucleotides, macromolecules) needed for rapid cell division. Therefore, increased tumor glucose uptake is typically a poor prognostic marker. We recently demonstrated in an analysis of PET-CT images from TCIA that body mass index (BMI) correlated negatively with the lean body mass-corrected maximum standard uptake value (SUVmax) in non-small cell lung cancer (NSCLC)^11^, consistent with the prior epidemiologic data. Yet, recent data in NSCLC has suggested that ^18^F-FDG does not correlate with glycolytic capacity per se, but more closely with proliferation index, begging both a deeper and a more comprehensive analysis of the metabolic pathways related to NSCLC nutrient metabolism^12^.

However, when pursuing the question of how excess body weight may be protective in lung cancer, it is important to recall that while most of those who fall into the clinically defined overweight and obese categories do so because of excess body fat, this is not universally the case. For example, athletes are often in the overweight or obese categories as designated by body weight, despite having low body fat and good metabolic health. In addition, certain populations have healthy range BMIs with poor lean muscle mass and excess relative adiposity. Consistent with this, previous analyses correlating anthropometric indices to visceral adipose tissue (VAT), a strong predictor of cardiometabolic risk, demonstrated that BMI did not correlate with VAT^9,10^. These data highlight the need to use markers of adiposity, rather than BMI, when examining how excess body weight affects cancer outcomes. To address this, we calculated skeletal muscle mass and visceral and subcutaneous abdominal adipose tissue from positron emission tomography-computed tomography (PET-CT) scans available in The Cancer Imaging Archive (TCIA). In addition, we measured tumor and tissue specific glucose uptake in the scans of the same patients to more comprehensively assess the metabolic activity of their tumors.

To assess elevated genes and biological activity with more granularity, we chose to extend our analyses to include RNA-sequencing and quantitation of tumor infiltrating leukocytes to obtain a deeper understanding of how body composition and tumor glucose uptake may influence the immunometabolic tumor microenvironment. In light of recent evidence that glucose uptake as detected on PET may reflect immune cell metabolism rather than tumor cells^13^, we examined tumor infiltrating leukocytes identified by caMicroscope and validated by pathologists in the tumors of patients with available PET/CT image data. In addition, RNA-sequencing has proven to reveal metabolic vulnerabilities through transcriptomic analyses of the tumor microenvironment^14^, so we employed TCGA analyses to gain a deeper understanding of the metabolic transcriptomic tumor landscape.

A unique aspect of the current study is the comparison between the two most common non-small cell lung cancer lung cancer types, LUAD and LUSC, which comprises 85% of all lung cancer cases. A recent analysis of data from more than 37,000 lung cancer patients found differences in demographics and in outcomes: LUSC patients tend to be older, more likely male, and more likely smokers^15^. Furthermore, first-line treatment protocols of these subtypes differ, and results regarding relative survival rates between these NSCLC subtypes vary, with improved survival in LUSC^16–19^ and LUAD^15,20–23^ having been reported. Further, recent genomic analyses suggest differences in immunogenicity between LUSC and LUAD.^24^

In this study, we employed a multimodal approach to understand how body composition, metabolism, and immune function intersects with tumor glucose uptake, and tumor genomics in NSCLC. We anticipate that these results could inform new precision medicine approaches to understand how body composition may alter tumor genomics and metabolism in lung cancer.

## Methods

Utilizing three complementary National Cancer Institute-funded open-source databases, The Cancer Imaging Archive (TCIA), The Cancer Genome Atlas (TCGA), and the Quantitative Imaging in Pathology (QuIP) with caMicroscope^25,26^, we performed a retrospective cross-sectional analysis to examine phenotypic, metabolic, and genomic intersections of adiposity and tumor metabolism in NSCLC. All patients with an ^18^F-FDG PET/CT scan available from TCIA were studied, and RNA-sequencing and histology was obtained for all subjects for whom this data was available. All subjects provided informed consent in accordance with each site’s institutional review board. Eighteen patients with lung adenocarcinoma and seventeen patients with squamous-cell carcinoma, both located in the bronchus and lungs were included.

### ^18^F-FDG PET/ CT Analysis

PET/CT images of TCGA-LUSC^27^ and TCGA-LUAD^28^ were accessed and downloaded via the TCIA Portal. One full-body CT and 2 PET images (1 attenuation corrected and 1 nonattenuation corrected-when available) were uploaded to the Image J platform using and open-source plug-in PET-CT Viewer, then co-registered and reconstructed.^29^ “Any” parameter to select any voxels meeting tissue density criteria of the PET-CT was used as described.^11,30,31^

SUVmax and SUVmean were obtained for tumor and normal tissues including brain, heart, liver, subcutaneous white adipose tissue (WAT), and skeletal muscle (deltoid). Tumor SUVmax was corrected to background ^18^F-FDG in the blood, and a tumor-to-descending aorta calculation was made for tumor SUV.

Volume of visceral, subcutaneous, and total adipose tissue was obtained with two consecutive CT slices between the L3 and L4 vertebrae with a −190 to −30 Hounsfield Unit cutoff.^31,32^ Skeletal muscle volume was obtained on the same slices by drawing regions of interest around all major abdominal muscles with a −29 to 50 Hounsfield Unit cutoff.^32^

### RNA-seq Expression Analyses

Differential expression analysis between LUAD and LUSC was conducted using processed data from XenaBrowser (https://xenabrowser.net; dataset: gene expression RNAseq – IlluminaHiSeq, dataset ID: TCGA.LUAD/LUSC.sampleMap/HiSeqV2, unit: log2(norm_count+1)).^33^ This public TCGA expression data was used to identify genes differentially expressed between high VAT subgroups in LUAD vs LUSC as well as high SubC groups in LUAD vs LUSC (P < 0.001; median VAT/SubC split). For TGFB pathway expression analysis, each gene of the ‘KEGG_TGF_BETA_SIGNALING_PATHWAY’ gene set (https://www.gsea-msigdb.org/gsea/msigdb/cards/KEGG_TGF_BETA_SIGNALING_PATHWAY) was used to assess the correlation using the scipy.stats python package. KEGG Gene Sets and GO Processes barplots presented were obtained using Enrichr (https://maayanlab.cloud/Enrichr/) using DEGs in LUAD vs LUSC.^34^ Gene sets shown are all significantly differentially enriched between groups (Adj. p <0.05). Expression heatmaps were generated using the ComplexHeatmap package in R.^35^

### Tumor Infiltrating Leukocyte Analyses

Tumor infiltrating leukocyte data was obtained from a convolutional neural network (CNN) applied to identify patches of cell subtypes, and were confirmed by a pathologist for accuracy.^36^ In brief, over 5,000 histological images from the TCGA database were computationally stained, and the CNN was trained with pathologist categorization of extracted patches. Features were extracted and proportions of individual cell types were quantified from the CNN analyses. In this analysis, fraction of the tumor that was TILs, and the subtype of cells were selected for comparisons among these studies.

### Statistics

In all comparisons between two groups, a student’s t test was performed to compare differences in means. Pearson r’s were calculated to examine relationships between two groups in GraphPad Prism Version 9.1. Two-tailed P values were computed. REMARK reporting guidelines were used where applicable.^37^ Statistical significance was determined as P values less than 0.05.

## Results

### Patient characteristics

We took a multi-modal approach to understand immunometabolomic interactions in LUSC and LUAD, analyzing datasets available in the National Cancer Institute-supported TCIA and TCGA databases. CT-defined visceral and subcutaneous adipose tissue and tissue-specific glucose uptake by ^18^FDG-PET, tumor RNA sequencing, and histological assessment of tumor-infiltrating leukocyte counts and subtypes were assessed (**Figure 1**). Comparing the two tumor types, patients were matched for age, weight, sex, race, and tumor stage (**Supplemental Table 1**), although subjects were limited to 17 and 18 per tumor type due to the limited numbers of patients for whom each of the aforementioned data sets were available in the TCIA and TCGA databases.

**Figure 1.**
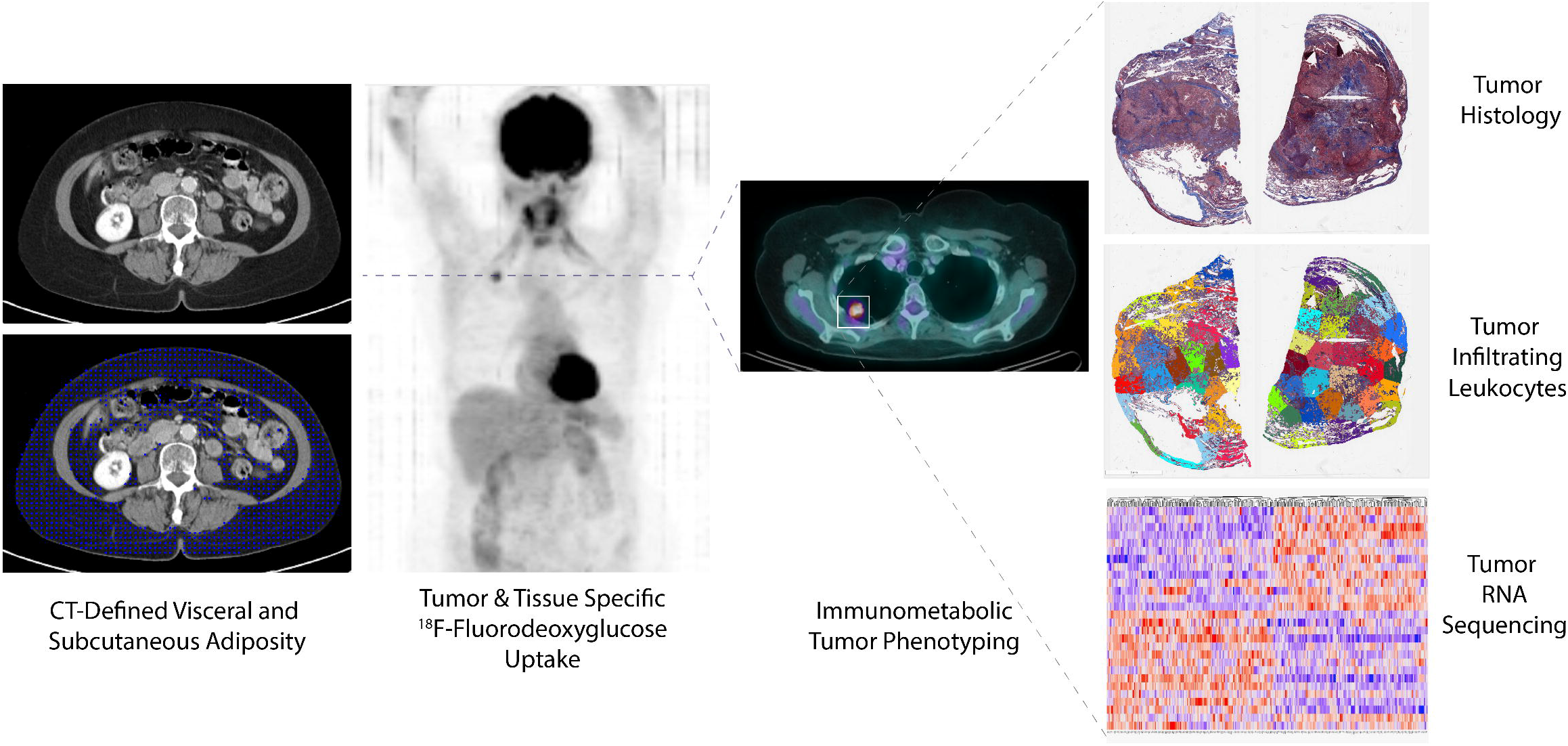
Overview of multimodal approach to immunometabolic phenotyping of Non-Small Cell Lung Cancer. 18F-FDG PET/CT images were analyzed to quantify visceral and subcutaneous adipose tissue and tissue specific glucose uptake. Within the tumors of NSCLC patients, we analyzed tumor infiltrating leukocyte profiles and RNA sequencing data all within the same cohort of patients. Data obtained from the TCGA and TCIA.

### Visceral and subcutaneous adiposity differentially affect tumor gene expression in NSCLC

We quantified visceral (VAT) and subcutaneous (SubQ) adipose tissue volume by PET-CT and found it to be unchanged between subjects with LUAD and LUSC tumors (**Figure 2A-B**), as in line with the identical body weight between patients with the two tumor types. Skeletal muscle volume was similarly unchanged between the tumor types (**Figure 2C-D**). However, high versus low (median split) adipose tissue mass was correlated with differences in gene expression in NSCLC tumors: VAT mass was associated with differences in expression of several Kyoto Encyclopedia of Genes and Genomes (KEGG) metabolic pathways involved in glucose, amino acid, and fatty acid metabolism, whereas Gene Ontology (GO) analysis revealed that pathways involved in neutrophil activation and function were differentially affected by VAT volume (**Figure 2E**). While KEGG pathway analysis painted a less clear picture of the physiologic relevance of the pathways enriched in association with SubQ adipose tissue volume, GO analysis again demonstrated differential expression of genes involved in neutrophil function and angiogenesis (**Figure 2F**).

**Figure 2.**
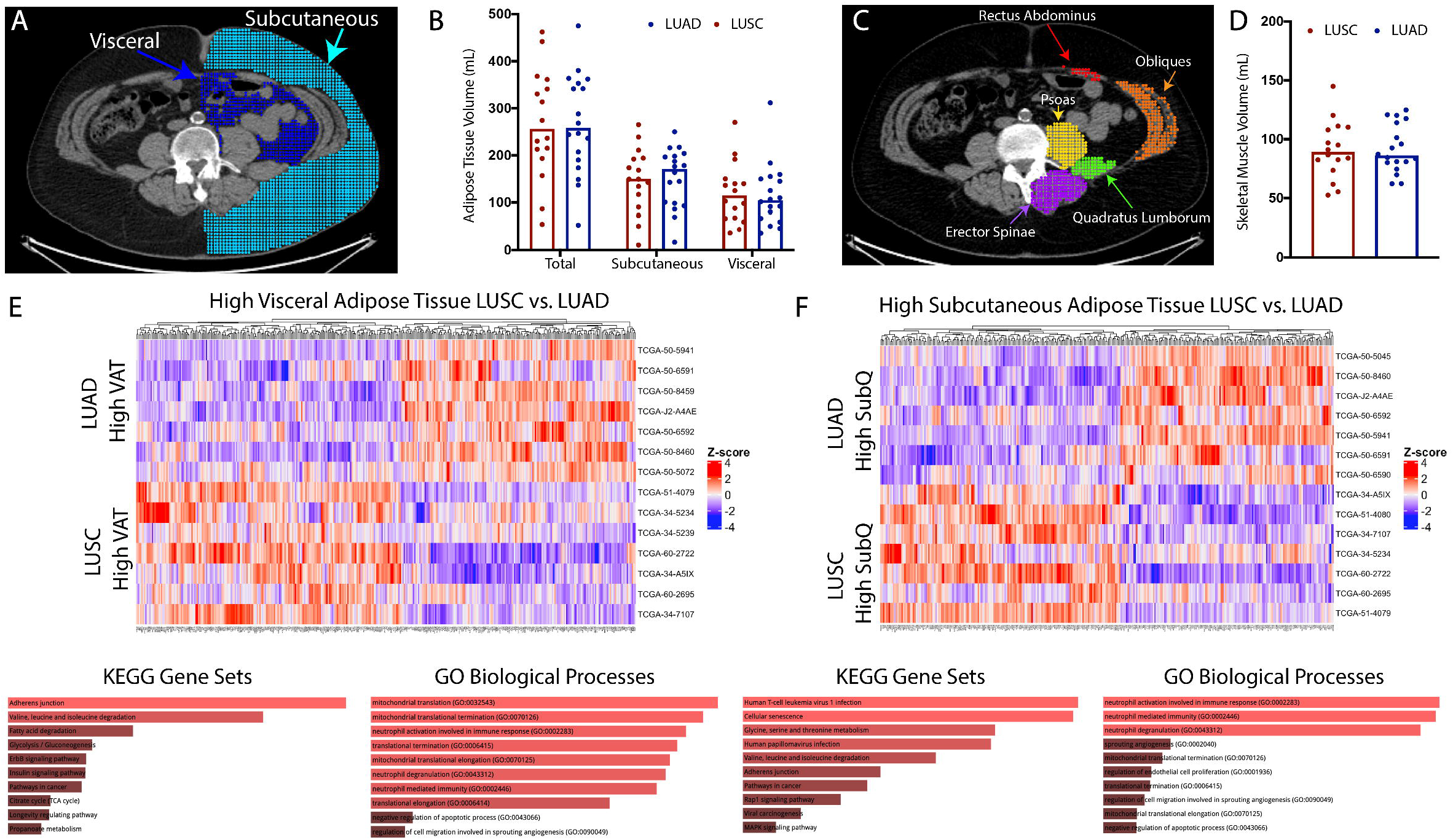
Visceral and subcutaneous adiposity differentially affect tumor gene expression in NSCLC. (A and B) visceral and subcutaneous adipose tissue was quantified from CT scans from the L3-L4 vertebral level in all patients. (C and D) abdominal skeletal muscle volume was quantified as the sum of five muscle groups at the L3-L4 vertebral level in all patients. Differential gene expression analyses with associated KEGG and GO processes for significantly differentially expressed (p<0.05) pathways was performed between LUAD and LUSC cohorts based on being in the top half of (E) VAT volume and (F) SubQ volume.

### The quantity and location of adipose tissue predicts differential gene expression patterns between LUAD and LUSC

Utilizing the TCIA database, we quantified tumor glucose uptake using ^18^F-FDG PET-CT and found that the maximum standardized uptake value (SUVmax) normalized to SUV in the descending aorta was higher in LUSC than in LUAD (p<0.0001) (**Figure 3A-C**), despite unchanged body weight, stage, or adipose tissue or muscle volume. Consistent with a tumor-specific effect rather than a systemic effect to promote increased glucose uptake, no differences were observed in ^18^F-FDG uptake in nontumor tissues including heart, liver, skeletal muscle (deltoid) adipose tissue (SubQ abdominal) and brain (**Figure 3D**). Having observed differences in glucose uptake without overall alterations in body composition between LUSC and LUAD, we aimed to determine whether body composition correlated with tumor glucose uptake within each tumor type. While VAT volume correlated positively with tumor glucose uptake in LUAD (p=0.0398), we observed a relatively inverse correlation between the proportion of abdominal fat at L3 that was located in the visceral (as opposed to subcutaneous) compartment in LUSC (p=0.07) (**Figure 3E-F**). In contrast, tumor glucose uptake did not correlate with SubQ adipose tissue volume in either tumor type (**Figure 3G**).

**Figure 3.**
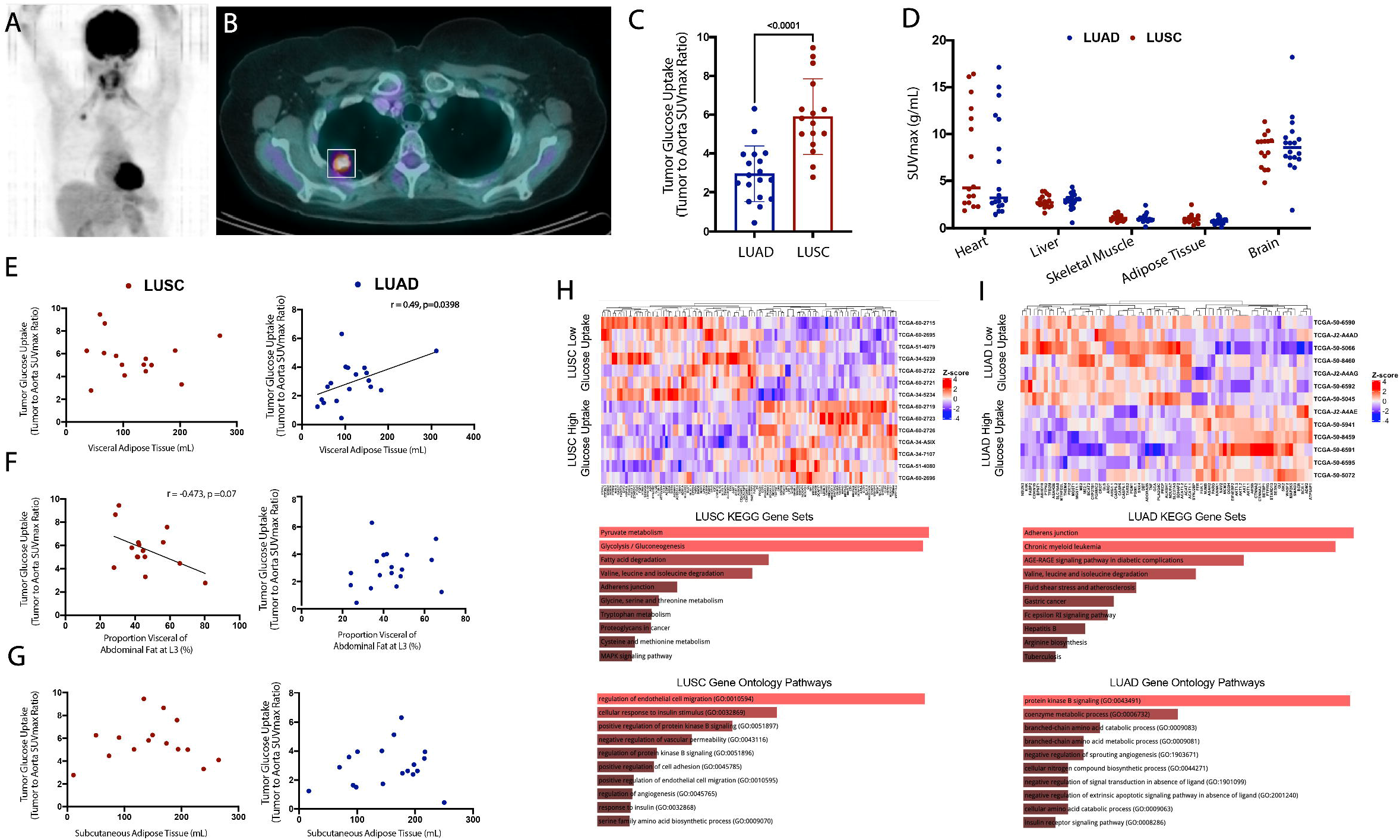
Tumor glucose uptake and metabolic genes within the tumor differ between LUAD and LUSC. (A) representative maximal intensity projection and (B) coronal PET/CT slice identifying the lung tumor. Tumor glucose uptake was quantified as a tumor to background ratio (C) and as SUVmax in healthy tissues (D). Correlation analyses between tumor glucose uptake and (E) absolute visceral adipose tissue volume, (F) proportion of adipose tissue volume that is visceral, and (G), absolute subcutaneous adipose tissue volume. Differential gene expression analyses with associated KEGG and GO processes for significantly differentially expressed (p<0.05) pathways was performed in (H) LUSC and (I) LUAD based on a median split within each cohort of ranked tumor glucose uptake values.

Next, we performed differential gene expression analysis on the tumors from NSCLC patients with high versus low tumor glucose uptake. This analysis revealed numerous differentially expressed KEGG metabolic pathways including pyruvate, glucose, amino acid, and fatty acid metabolism, as well as GO biological processes including angiogenesis and insulin signaling processes in LUSC (**Figure 3H**). In contrast, LUAD gene expression showed a less clear association between tumor glucose uptake and metabolic gene expression, while expression of genes in angiogenesis GO pathways did correlate with tumor glucose uptake in LUAD (**Figure 3I**).

### Tumor infiltrating leukocytes correlate with tumor glucose uptake in both NSCLC tumor types

The TCGA database provides unique access to data on tumor infiltrating leukocyte (TIL) density and cell type.^36^ Although we did not observe any differences in TIL number or subtype, not surprisingly, we observed a negative correlation between TIL fraction and tumor glucose uptake in both LUAD (p=0.017) and a trend towards an association in LUSC (p=0.16) (**Figure 4A-E**), consistent with a tumor-suppressive effect of TIL infiltration. Further, there was a trend towards a negative correlation between TIL content and VAT content in the LUSC cohort (p=0.08).

**Figure 4.**
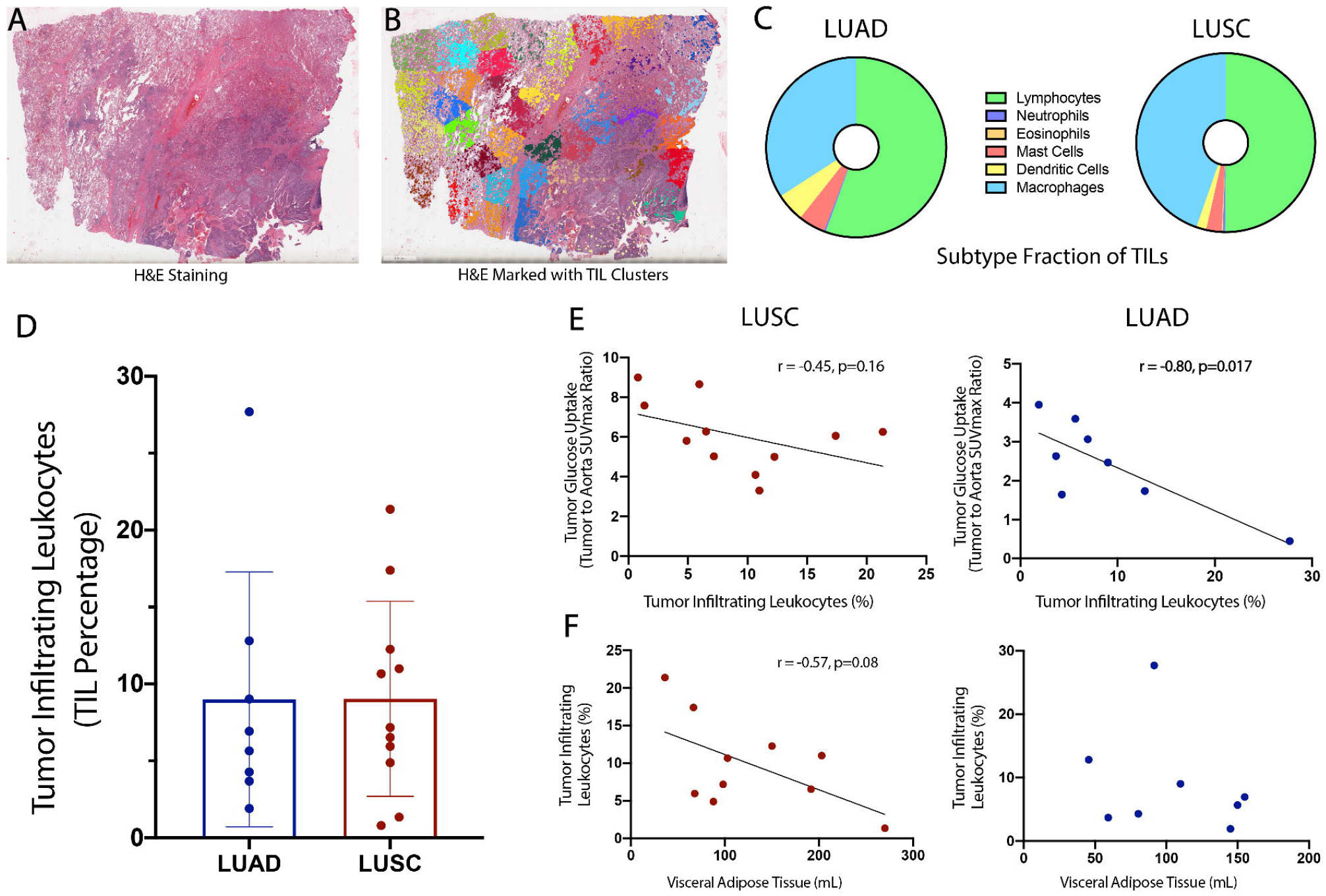
Tumor infiltrating leukocytes are negatively correlated with tumor glucose uptake and visceral fat content in LUSC and LUAD. A representative histological tumor slice with (A) H&E only, or (B) clusters identifying different immune cell populations within the tumor slide. (C) the subtyping of the tumor infiltrating leukocytes (TILs) in LUAD and LUSC tumors. (D) proportion of tumor cells that were identified as TILs in both cohorts. Correlation analyses between TIL fraction and (E) tumor glucose uptake and (F) visceral adipose tissue volume.

### TGF-β2 expression correlates differentially with visceral fat in LUSC and LUAD

Transforming Growth Factor-β (TGF-β) was identified to be previously implicated in excess adiposity^38,39^, immune function^40^, and cancer progression^41^. To examine possible biological mediators of the visceral adiposity’s influence on the tumor microenvironment, we performed correlational analyses between enriched genes and visceral adipose tissue content. Of all genes in the KEGG TGF-β pathway, TGF-β2 was most significantly related to VAT in LUSC. We observed differential correlations between TGF-β2 expression and VAT in the two tumor types: visceral fat volume was positively correlated with tumor TGF-β2 expression (p=0.0006) in LUSC, but unrelated to tumor TGF-β2 expression in LUAD (**Supplemental Figure 1**). These data suggest that the impact of systemic metabolism on tumor TGF-β2 signaling deserves further attention in NSCLC, and highlights the importance of differentiating between NSCLC tumor types when investigating the impact of this pathway in lung cancer.

## Discussion

The relationship between systemic metabolism and subtypes of lung cancer is undoubtedly complicated. Although it is clear that increased tumor SUVmax predicts poor outcomes in lung cancer^42–45^, how systemic metabolism and, specifically, adiposity affects tumor glucose metabolism remains unclear. It appears that overweight and obesity improve the response to immunotherapy in lung cancer^7,8^; however, it is not known whether obesity confers improvements in prognosis in all types of lung cancer, or if this effect is tumor type-specific. In addition, the use of the commonly used and clinically accessible parameter BMI to predict cancer outcomes has come under scrutiny. This is in part because while BMI is tightly correlated with body fat at the population level, it is a poor predictor of adipose tissue mass at the individual level^46,47^. Total body fat mass is a better predictor of cardiometabolic health^47–50^ and cancer risk^51,52^ than BMI. As subcutaneous adipose tissue is generally considered metabolically inert and far less of a cardiometabolic risk factor than visceral and ectopic adipose tissue^53–55^, both the quantity and the location of adipose tissue may be predictive of cancer outcome. Consistent with a tumor type-dependent effect of systemic metabolism to modulate lung cancer initiation and/or progression, patients with non-alcoholic fatty liver disease (i.e. increased ectopic lipid content) were more likely to have non-squamous cell carcinoma^56^.

Therefore, in this study, we performed a multi-modal analysis of the relationship between body composition, tumor glucose uptake, immune cell infiltration and subtypes, and tumor genomic in the two major subtypes of NSCLC, LUAD and LUSC. Surprisingly, we identified opposite correlations between visceral adipose tissue volume and tumor glucose uptake in the two tumor types: VAT correlated positively with SUVmax in LUAD and strongly tended to correlate negatively with SUVmax in LUSC (**Figure 3E-F**). These data are consistent with a recent meta-analysis demonstrating that insulin signaling and fatty acid metabolism pathways, which tend to be downregulated in obesity, are enriched in LUSC but unchanged in LUAD^57^, suggesting a protective effect of obesity acting through these pathways in LUSC. This is consistent with both epidemiologic data^7,8^ and with our recent PET-CT analysis^11^.

Both the SUVmax and RNA expression data obtained in this study are consistent with a greater influence of metabolism on tumor progression in LUSC as compared to LUAD. Despite unchanged tumor stage, adiposity, and non-tumor tissue glucose uptake, tumor SUVmax in LUSC was twice that of LUAD. Differentially expressed KEGG pathways in tumors of LUSC patients with high versus low tumor glucose uptake were enriched for metabolic pathways, including glucose, pyruvate, fatty acid, and amino acid metabolism, whereas LUAD KEGG analysis provided a less compelling argument for a critical role of regulation of tumor metabolism (**Figure 3H-I**). These data hint that metabolic adjunct approaches may be more likely to be effective in LUSC as compared to LUAD, and should consider the fact that glucose is likely not the only substrate that contributes to tumor progression in these tumor types.

As NSCLC can be a highly immunogenic tumor type responsive to immunotherapy, it is important to consider how systemic metabolism and body composition may intersect with tumor immune cell infiltration. The fraction of immune cell subtypes may correlate with tumor stage in NSCLC: Zhang et al. demonstrated decreases in the fraction of M0 macrophages and memory B cells in advanced stage as compared to early stage LUAD, but no difference in the fraction of immune cell subtypes in advanced stage as compared to early stage LUSC^58^. In the current study, increased TILs correlated with lower glucose uptake, predicting improved prognosis, but neither TIL number nor subtype differed between LUSC and LUAD. These data suggest that tumor leukocyte infiltration is a metabolism-independent predictor of prognosis in NSCLC, and may posit that the immune cell fraction of a tumor is not the dominant glucose consumer in LUAD, and possibly LUSC.

However, although there was no clear association of TILs themselves with NSCLC subtype, this should not be taken as an indication that proteins downstream of TILs may not be associated with systemic metabolism or body composition in a tumor typespecific manner. Interestingly, we observed a positive relationship between visceral fat volume and *Tgfb2* gene expression in LUSC, but no relationship in LUAD (**Supplemental Figure 1**). TGF-β2 is a cytokine that is produced by a variety of cell types, including lymphocytes, monocytes/macrophages, and many parenchymal cells, and it appears to play a complicated role in cancer: in early stages, it limits cell division by inhibiting cell cycle progression via inhibition of cyclin expression^59^; however, in late stages, when mechanisms of cell cycle regulation have already been overcome, it promotes induction of the epithelial to mesenchymal transition^60^, thereby enhancing cell migration and metastasis. However, because in our study there was no difference in tumor stage between subjects with LUAD and LUSC, the differing relationship between VAT and *Tgfb2* expression cannot be attributable to discrepancies in tumor stage. However, these results raise the intriguing possibility that TGF-β2 may be a metabolism-responsive biomarker which may have differential implications for prognosis in LUAD and LUSC. In addition, TGF inhibitors have been used clinically, and it should be further examined whether TGF-β2 inhibition could play a role in tumor suppression in LUSC patients with visceral adiposity.

In summary, here we performed a comprehensive analysis of the impact of body composition on human tumor glucose metabolism, transcriptomic landscape, and immune cell infiltration and subtypes using the publicly available NCI-supported TCIA and TCGA databases. Subcutaneous and visceral fat content is differentially related to the tumor metabolic landscape in LUAD vs. LUSC tumor types. In addition to having higher tumor glucose uptake, the LUSC cohort had a much greater enrichment of metabolic genes, suggesting that nutrient metabolism (not only glucose, but fatty acids and amino acids as well) plays a dominant role in the progression of LUSC, and to a lesser degree in LUAD. Further, our clinical data suggest that higher TILs did not have a positive association with tumor glucose uptake in either NSCLC subtype, in contrast to findings from recent animal model data.^13^ In this study, we ultimately provide insight into the differential immunometabolic physiology in LUAD and LUSC tumor types, and suggest that adjunctive metabolic cancer therapy may be a more promising approach in LUSC.

## Acknowledgments

This study was funded by grants from the U.S. Public Health Service (K99/R00 CA215315 and a Career Enhancement Program Award from the Yale SPORE in Lung Cancer [1P50CA196530] to R.J.P. B.P.L and K.B.G. are supported under the National Institutes of Health Medical Scientist Training Program Training Grant T32GM007205. KP was also supported by the Yale SPORE in Lung Cancer [1P50CA196530].

---

**Supplemental Figure 1.** TGFB2 is positively correlated with VAT in LUSC but not LUAD. Correlations were performed between visceral fat volume and TGFB2 mRNA expression in the tumor of (A) LUSC and (B) LUAD.

**Supplemental Table 1.** Patient demographics for LUAD and LUSC cohorts. N/A indicates that the data were not available.

