## Supplementary figures and images for "Multimodal Analysis Reveals Differential Immuno-Metabolic Features in Lung Squamous Cell Carcinoma and Adenocarcinoma"

### Supplemental Figure 1

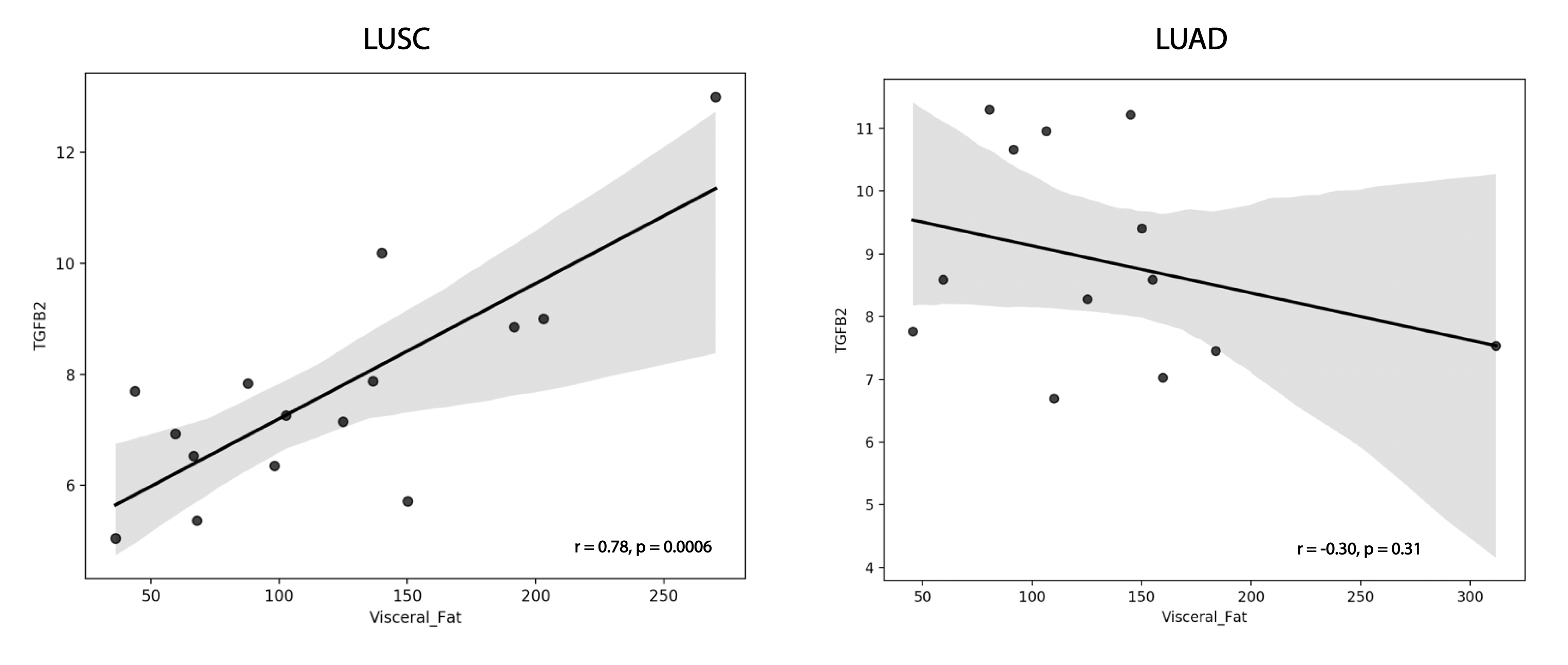
