## Supplemental Table 1 for "Multimodal Analysis Reveals Differential Immuno-Metabolic Features in Lung Squamous Cell Carcinoma and Adenocarcinoma"

|  |  | LUAD (n=18) | LUSC (n=17) |
| --- | --- | --- | --- |
|  |  | Mean ± SD | Mean ± SD |
| Age (years) |  | 67.7 + 6.3 | 70.9 + 8.6 |
| Sex | Male | 8 | 10 |
|  | Female | 9 | 7 |
|  | N/A | 1 | 0 |
| Body Weight (kg) |  | 71.2 + 13.1 | 75.6 + 15.2 |
| Stage | I | 6 | 7 |
|  | II | 1 | 8 |
|  | III | 4 | 2 |
|  | IV | 1 | 0 |
|  | N/A | 6 | 0 |
| Race | Black | 2 | 4 |
|  | White | 10 | 13 |
|  | N/A | 6 | 0 |
